# Physically active lifestyle is associated with attenuation of hippocampal dysfunction in healthy older adults

**DOI:** 10.1101/2021.05.27.445934

**Authors:** Tamir Eisenstein, Nir Giladi, Talma Hendler, Ofer Havakuk, Yulia Lerner

## Abstract

Alterations in hippocampal function have been shown in older adults, expressed as changes in hippocampal activity and connectivity. While hippocampal activation during memory demands have been demonstrated to decrease with age, some older individuals present increased activity, or hyperactivity, of the hippocampus which is associated with increased neuropathology and poorer memory function. In addition, lower functional coherence between the hippocampus and core hubs of the default mode network (DMN), namely the posteromedial and medial prefrontal cortices, as well as increased local intrahippocampal connectivity, were also demonstrated in cognitively intact older adults. Aerobic exercise has been shown to elicit neuroprotective effects on hippocampal structure and vasculature in aging, and improvements in maximal aerobic capacity (MAC) have been suggested to mediate these exercise-related effects. However, how these lifestyle factors relate to hippocampal function is not clear. Fifty-two cognitively intact older adults (age 65-80) have been recruited and divided into physically active (n=29) or non-active (n=23) groups based on their aerobic activity lifestyle habits. Participants underwent resting-state as well as task-based fMRI experiments which included an associative memory encoding paradigm followed by a post-scan memory recognition test. In addition, forty-four participants also performed cardiopulmonary exercise tests to evaluate MAC. While both groups demonstrated increased anterior hippocampal activation during memory encoding, physically active lifestyle was associated with significantly lower activity level and higher memory performance in the recognition task. In addition, the physically active group also demonstrated higher functional connectivity of the anterior and posterior hippocampi with the core hubs of the DMN, and lower local intra-hippocampal connectivity within and between hemispheres. MAC was negatively associated with hippocampal activation level and demonstrated positive correlation with hippocampal-DMN connectivity. According to these findings, aerobically active lifestyle may be associated with attenuation of hippocampal dysfunction in cognitively healthy older adults.

## 1. Introduction

Episodic memory, the ability to encode, consolidate, and retrieve past experiences and events, is one of the most affected cognitive abilities during aging ^1^. The hippocampus is a brain region which plays a key role in episodic memory processes ^2–4^, through anatomical and functional interactions with cortical and sub-cortical regions across the brain ^5,6^. The hippocampus presents distinct structural and functional connections along its longitudinal axis with its anterior part being more connected to anterior temporal and ventromedial prefrontal regions, and its posterior part more linked to posterior-medial cortices and hubs of the default mode network (DMN) such as the posterior cingulate cortex ^6^. Moreover, from a cognitive perspective, the anterior hippocampus has been shown to be more associated with memory encoding while its posterior part has been more related to retrieval ^7^.

Older adults have been demonstrated to exhibit age-related alterations in hippocampal function. Hippocampal activity patterns during memory demands have been shown to generally decrease with age ^8–10^. Furthermore, this age-related hypoactivity patterns was suggested to be region and process selective, affecting more pronouncedly the anterior part of the hippocampus during encoding ^8,10^. However, some older individuals demonstrate abnormally increased hippocampal activation during memory encoding and retrieval compared to both old and younger counterparts, also referred to as hippocampal hyperactivity. In addition, although this phenomenon has been observed in both healthy aging and prodromal stages of Alzheimer’s disease (AD), such as mild cognitive impairment (MCI), it is not clear whether this activity pattern represents a compensatory effect, or a dysfunctional pathological consequence.

While increased hippocampal activity or hyperactivity has been generally demonstrated to be associated with poorer neurobiological and cognitive outcomes in cognitively healthy older adults, it has been both negatively and positively linked with cognitive and clinical outcomes in patients with MCI ^11–15^. For example, increased left hippocampal activation during memory encoding in MCI was correlated with better memory performance and cognitive-clinical status in some studies ^11,12^, while others found it to be associated with poorer memory performance and increased Aβ burden ^14,15^.

In addition to altered activation levels, changes in the resting state functional connectivity (RSFC) of the hippocampus have also been documented to take place during aging. The hippocampus constitutes a part of the medial temporal subsystem of the DMN and is functionally and anatomically connected with regions comprising this network ^16^. However, evidence suggests that with increasing age hippocampal activity becomes less coherent with the activity of distant brain regions such as major hubs of the DMN including the posteromedial and medial prefrontal cortices (or decreased HC-DMN RSFC) and more locally connected within itself (or increased intra-HC RSFC) ^17–19^. Furthermore, this increased local connectivity was found to be associated with increased AD pathology burden in distinct memory networks and poorer episodic memory performance.

While neurobiological alterations have been well documented in the hippocampus during aging, several lifestyle factors have been proposed to attenuate this age-related deterioration, one of which is physical activity. Physically active lifestyle has been associated with prevention of cognitive decline and risk of AD and dementia ^20–22^. Aerobic exercise has been associated with hippocampal neuroprotective effects in both animal models ^23–27^, and healthy human subjects ^28–30^. However, studies examining the effect of physical exercise on the hippocampus in older adults had largely focused on structural characteristics and to our knowledge no previous work had focused specifically on the relationship between aerobically active lifestyle and hippocampal function in human aging.

Several factors have been suggested to potentially mediate the effects of aerobic exercise on the brain. Maximal oxygen consumption is perhaps the most studied physiological correlate of aerobic fitness in this context. It reflects the body’s peak metabolic rate in generating adenosine triphosphate (ATP) molecules through aerobic metabolism, or maximal aerobic capacity (MAC) ^31^. Previous studies that demonstrated neuroprotective effects of aerobic exercise intervention in older adults also found favourable structural and cerebrovascular hippocampal changes to correlate with improved MAC ^28–30^. However, no study to date to our knowledge specifically examined the relationship between MAC and age-related hippocampal dysfunction patterns.

The current study aimed to address this gap by investigating the relationship between lifestyle’s aerobic exercise and hippocampal function during both resting state and active memory demands in cognitively healthy older adults. In addition, we aimed to examine whether the extent of these trends may be associated with the level of MAC. We first aimed to examine the activity levels of the anterior hippocampus during associative memory encoding task, since this hippocampal subpart has been shown to be more involved in memory encoding processing ^7^. Associative memory paradigm was chosen as this form of episodic memory has been shown to be highly sensitive to aging compared to item memory, and relies greatly on hippocampal function ^32^. Next, we aimed to investigate both remote and local RSFC patterns of both anterior and posterior hippocampal subparts. Distant hippocampal RSFC was examined with the two core hubs of the DMN, i.e., the posteromedial and medial prefrontal cortices (HC-DMN RSFC), and intra-HC RSFC patterns were examined between bilateral anterior and posterior hippocampi. We then examined whether physically active individuals may demonstrate distinct patterns of hippocampal activity and RSFC compared to sedentary individuals, and whether it may explain differences in memory performance between the two groups. Lastly, we investigated the correlations between MAC and all hippocampal and memory measures. According to the previous findings in the literature, we hypothesized that physically active older adults will demonstrate lower anterior hippocampal activity during memory encoding and higher memory performance, higher distant functional connectivity with the cortical DMN hubs and lower within hippocampal functional correlation. In accordance, we hypothesized that MAC, as a potential mediator of aerobic exercise, will demonstrate dose-response correlations with hippocampal function and memory measures in the same directions as physically active lifestyle.

## 2. Materials and methods

### 2.1 Participants

Fifty-two older adults aged 65-80 were recruited for the study (22 women/30 men). All participants were recruited from the community, were fluent Hebrew speakers, and reported no current or previous neuropsychiatric disorders (including subjective cognitive/memory decline or depressive symptoms) or any other current significant unbalanced medical illness (e.g., cardiovascular or metabolic). The research was approved by the Human Studies Committee of Tel Aviv Sourasky Medical Center (TASMC), and all participants provided written informed consent to participate in the study.

### 2.2 Aerobic activity lifestyle habits assessment

Participants’ current aerobically active lifestyle was assessed using a background interview that examined background characteristics as well as the physical exercise habits of the participants during the last year. The interview was administered face-to-face on the first assessment day. Specifically, participants were asked regarding the amount of weekly exercise-oriented activity sessions of some type of aerobic exercise (i.e., walking, running, cycling, swimming, elliptical/cross training gym machines, etc..) or *“How often during the week did you participate in leisure time aerobic activity for the purpose of sporting or exercise that lasts at least 20 min in the passing year?”*. Next, based on their reported activity patterns participants were divided into two groups, active (n=29) and non-active (n=23). Frequency of twice-a-week or higher was set as the cut-off for active lifestyle based on previous methodologies ^22,33^, findings ^34^, and recommendations ^35,36^. In addition, since we used a subjective measure of physical activity, using a cut-off of several times-per-week may help to decrease potential information bias from participants’ reports on their true activity levels. Also, regarding the potential of self-selection bias of volunteering to the study, it is important to note that the fraction of the active group out of the whole sample (56%) was similar to a recent report on aerobically active lifestyle habits among older adults in the areal county from which participants were recruited (53.5% who reported being active) ^37^.

### 2.3 Maximal aerobic capacity measurement

Forty-four participants (16 women/28 men) of the total 52 participants also underwent a graded maximal cardiopulmonary exercise test performed on a cycle ergometer to evaluate MAC. Assessments were conducted at the Non-Invasive Cardiology Outpatient Clinic at TASMC. Tests were supervised by a cardiologist and an exercise physiologist, while continuously monitoring for cardiopulmonary parameters, including oxygen consumption (VO2), heart rate, blood pressure, and respiratory exchange ratio (RER). An automated computerized ramp protocol was used to increase exercise intensity by 10 Watt/minute for women and 15 Watt/min for men, while participants were asked to maintain a constant velocity of 60 revolutions per minute. The highest average VO2 value recorded during an intensity interval (2/2.5 watts increment for women and men, respectively) was considered as the MAC value obtained from the procedure and that was used in further analyses.

### 2.4 Experimental procedure and stimuli

#### 2.4.1 Inside the MRI scanner

A face-name associative memory encoding task based on the classic paradigm by Sperling et al. ^38,39^ was used to evaluate hippocampal function. During scanning, participants were shown novel images of non-famous faces, each face paired with a fictional name (see **Figure 1A**). Participants were asked to memorize which name was coupled with each face, and to subjectively decide (by pressing a button) whether the name “fits” or not to the face. This subjective decision has been shown to enhance associative encoding ^38^. Participants performed one run of a block-design picture-viewing paradigm consisted of 8 blocks with 4 faces presented in each block (32 faces overall). Between blocks and consecutive in-block images participants were shown “resting” fixation blocks of a white cross in the middle of a black background. Prior to the scanning session, participants underwent a familiarization practice with the task to minimize novel task learning effects which could bias the neurocognitive outcome. Each stimuli block lasted for 21 seconds, while the overall run lasted 5:06 minutes.

**Figure 1.**
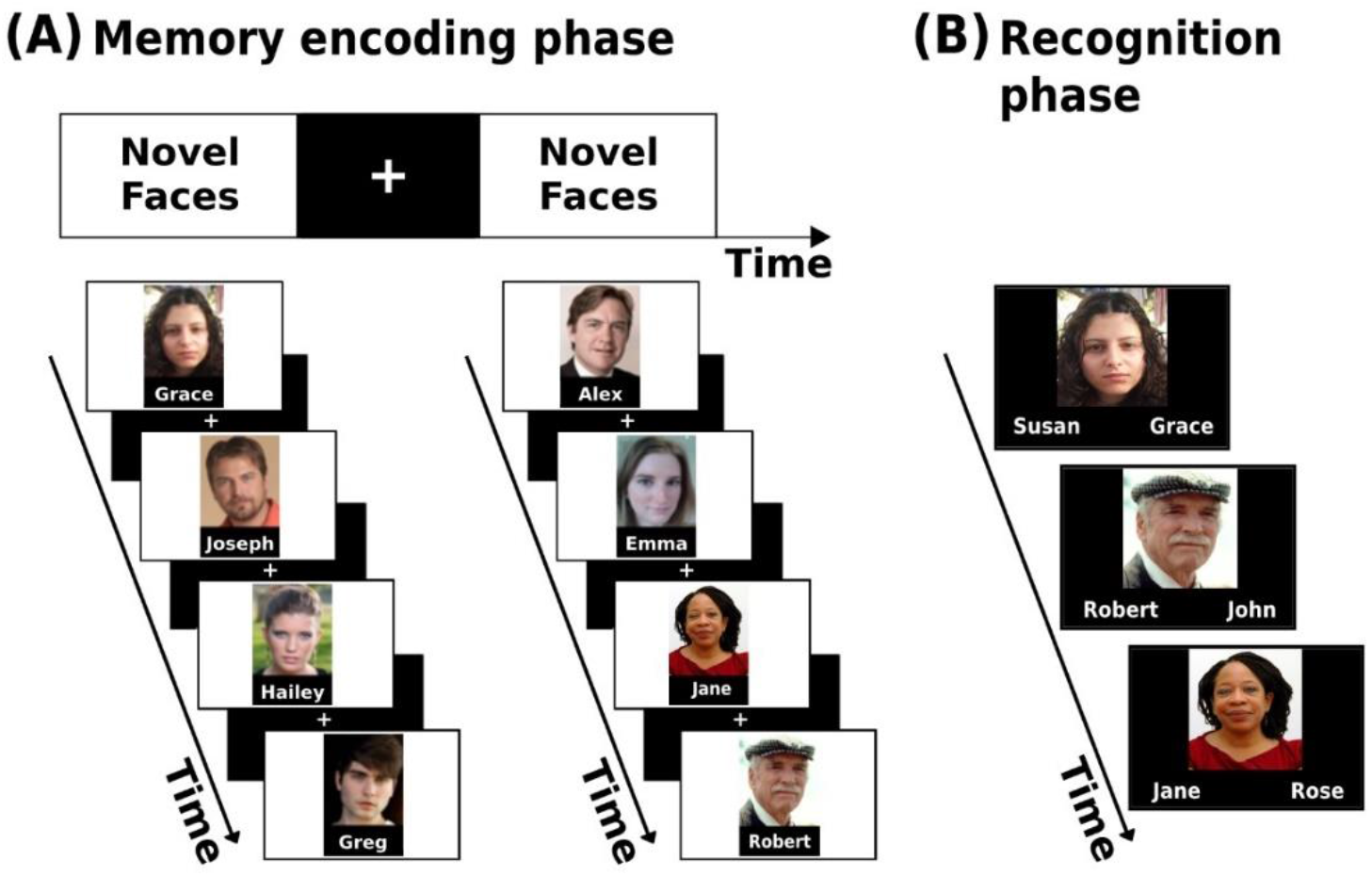
Associative memory encoding paradigm conducted in the scanner (A) and the post-scan memory recognition task (B).

#### 2.4.2 Outside the scanner – memory performance evaluation

Following the acquisition phase in the scanner, participants performed a two-alternative forced choice recognition task. During the task participants were shown images of all 32 faces presented during the encoding task and were asked to decide between two options which name was paired with each face (**Figure 1B**). The scores on the task were later used as the measure of memory performance.

### 2.5 MRI data acquisition

MRI scanning was performed at TASMC on a 3 T Siemens system (MAGNETOM Prisma, Germany). High resolution, anatomical T1-weighted images (voxel size = 1 ⨯ 1 ⨯ 1 mm) were acquired with a magnetization prepared rapid acquisition gradient-echo protocol with 176 contiguous slices using the following parameters: field of view (FOV) = 256 mm; matrix size = 256 × 192; repetition time (TR) = 1740 ms; echo time (TE) = 2.74 ms, inversion time (TI) = 976 ms, flip angle (FA) = 8°. These anatomical volumes were used for structural segmentation and co-registration with functional images. Blood oxygenation level dependent (BOLD) functional MRI was acquired with T2*-weighted imaging. The memory encoding task was conducted using the following parameters: 102 TRs of 3000 ms each; TE = 35 ms; FA = 90°; FOV = 220 mm; matrix size = 96 ⨯ 96; 44 slices, size = 2.3 ⨯ 2.3 ⨯ 3 mm, no gap (5:06 min run). The resting state run was carried out with the same parameters with 120 TR’s (6:00 min run). To minimize head movements, participants’ head were stabilized with foam padding. MRI-compatible headphones (OPTOACTIVEtm) were used to considerably attenuate the scanner noise and communicate with the participants during the session. Designated software (Presentation®, Neurobehavioral Systems) was used for visual stimuli presentation.

### 2.6 Functional MRI analysis

#### 2.6.1 Hippocampal activation

FMRI analysis was carried out using the FEAT tool in FSL 6.00 (FMRIB’s Software Library, www.fmrib.ox.ac.uk/fsl). The first 5 TRs of the functional data were discarded to allow steady-state magnetization. Registration of the functional data to the high resolution structural images was carried out using boundary based registration algorithm ^40^. Registration of high resolution structural to standard space (MNI152) was carried out using FLIRT ^41,42^, and then further refined using FNIRT nonlinear registration. Motion correction of functional data was carried out using MCFLIRT ^42^, non-brain removal using BET ^43^, spatial smoothing using a Gaussian kernel of FWHM 5 mm, grand-mean intensity normalisation of the entire 4D dataset by a single multiplicative factor, high-pass temporal filtering was performed with a Gaussian-weighted least-squares straight line fitting with a cut-off period of 100s. Time-series statistical analysis (pre-whitening) was carried out using FILM with local autocorrelation correction ^44^. A first-level task regressor of interest was defined and convolved using the blocks onset times with a double-gamma hemodynamic response function as well as a temporal derivative regressor of the task timing. In addition, 24 nuisance motion regressors were added to each first-level model and included 6 standard motion parameters (3 rotations, 3 translations), their temporal derivatives, and squares of all the above. Moreover, volumes with excessive head motion (predetermined as framewise-displacement value>0.9 mm) were scrubbed by adding an additional regressor for each volume to be removed. participants were removed and excluded from the group analysis if they had 30% or more of their volumes scrubbed out (one subject from the non-active group). Z statistic images were thresholded non-parametrically using clusters determined by Z>3.1 and a corrected cluster significance threshold of *p*=.05. Group-level analysis was carried out using FLAME (FMRIB’s Local Analysis of Mixed Effects) stage 1 ^45–47^. Group-level Z statistic images were thresholded non-parametrically using clusters determined by Z>2.3 and a corrected cluster significance threshold of *p*=.05. One other participant from the non-active group did not go through the encoding run. The mean activation patterns observed during the encoding task of the entire sample were used to identify anterior hippocampal regions which demonstrated increased activity during the task. Then, a mask was created for these regions (for each hemisphere) by masking the activation map with anatomical anterior hippocampal masks from (https://neurovault.org/collections/3731/) based on ^48^ (**Figure 2A**). Then, we used the FSL’s *Featquery* tool which enables the interrogation of FEAT results within a specific mask, to extract the parameter estimates from the bilateral anterior hippocampal regions activated during the task. The parameter estimate values were then used to examine differences in activation-level between the groups.

**Figure 2.**
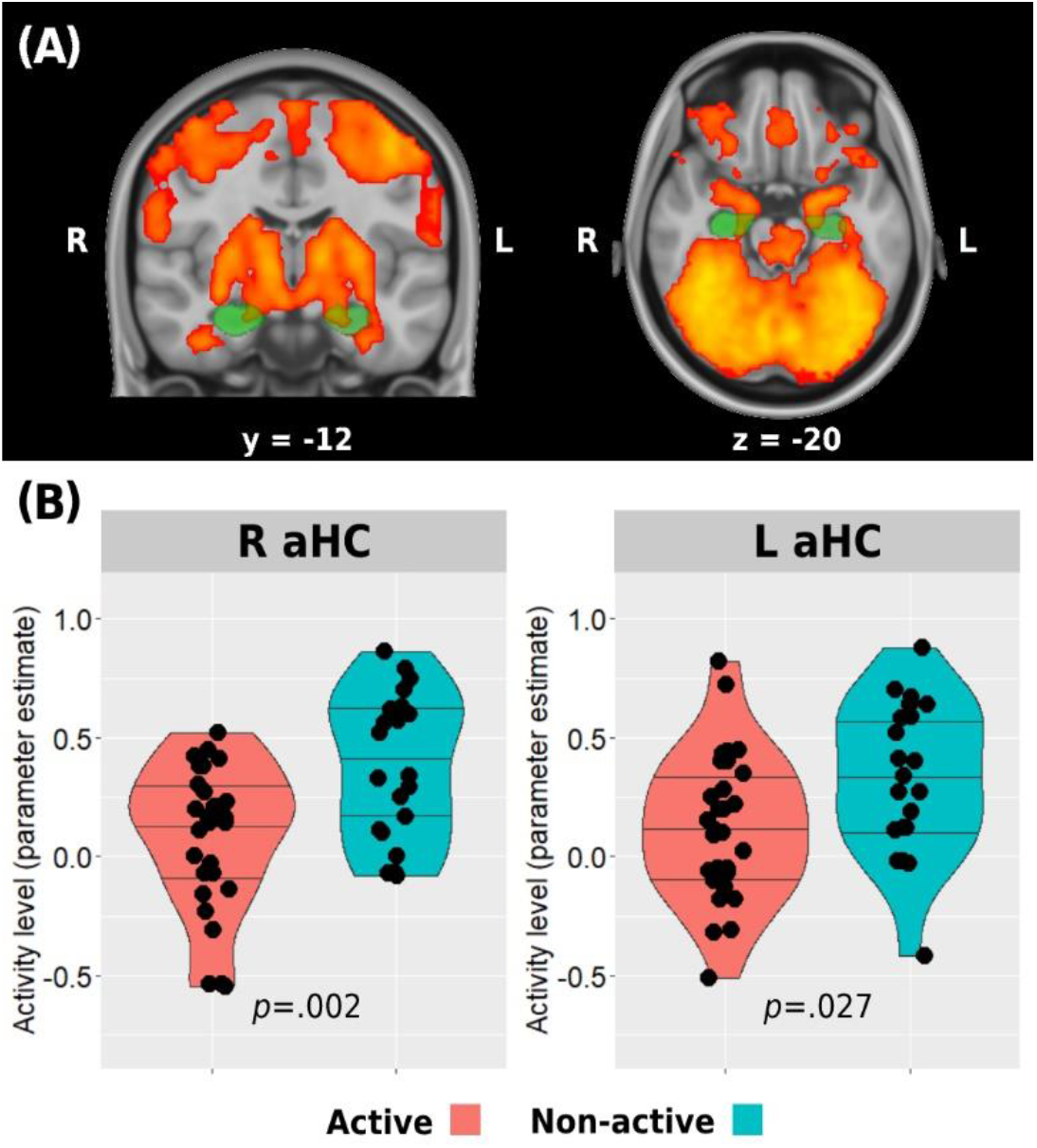
Anterior hippocampus activation during memory encoding. Whole sample mean activation during the task is presented in (A). Green regions represent anatomical masks of anterior hippocampi, while activated voxels within these areas were used to extract activity level for between-group analysis presented in (B). Horizontal lines in between-group plots represent 25%, 50%, and 75% quantiles. aHC = anterior hippocampus; L = left; R = right

#### 2.6.2 Hippocampal RSFC

Resting-state functional connectivity was carried out using the functional cconnectivity toolbox *CONN* v.19c (nitrc.org/projects/conn). Preprocessing included discarding the first 5 TRs to allow steady-state magnetization. Functional images were slice-time corrected, realigned to the middle volume, motion corrected, and normalized to the standard MNI152 space. Spatial smoothing was performed using a 5 mm FWHM Gaussian kernel. In order to reduce noise, functional volumes were bandpass filtered at 0.008–0.15 and component-based method (CompCor) was used to extract noise signals (e.g., white matter, CSF, and movement artifact) that were used as nuisance regressors to denoise the data. In addition, images that were regarded as movement outliers were regressed out. Movement outlier volumes were detected using the ART toolbox (nitrc.org/projects/artifact_detect/) and defined as volumes with a movement greater than 0.9 mm or signal intensity changes greater than 5 SD. These volumes were also used as nuisance regressors at the denoising step. No participant demonstrated more than 30% removed volumes. One participant from the active group did not go through the resting-state run. Remote functional connectivity of the anterior and posterior hippocampus with the main hubs of the DMN was examined by creating specific masks of these regions of interest. Anterior and posterior hippocampal masks were again created from (https://neurovault.org/collections/3731/) (**Figure 4A**). To create the DMN masks we used a deactivation contrast in the memory encoding task to elicit areas demonstrating decreased activity during the task (**Table S1**). The two main and larger clusters revealed from this analysis were anatomically corresponding to the posteromedial and medial prefrontal cortices and were used to create the masks for the HC-DMN RSFC analyses. The mean z-transformed correlation value between each pair of these eight areas were then used to examine differences in hippocampal-DMN functional connectivity between the groups. Intra-hippocampal resting state connectivity was examined in two ways. First, computing the z-transformed correlation between the time-series of bilateral anterior hippocampi and bilateral posterior hippocampi. Second, we calculated the local correlation within each hemispheric anterior or posterior regions using the Local Correlation implemented in *CONN*. This index represents a measure of local coherence at each voxel, characterized by the strength and sign of connectivity between a given voxel and the neighbouring regions in the brain ^49^. We used a 6 mm kernel for the Gaussian weighting function characterizing the size of the local neighbourhoods.

**Figure 3.**
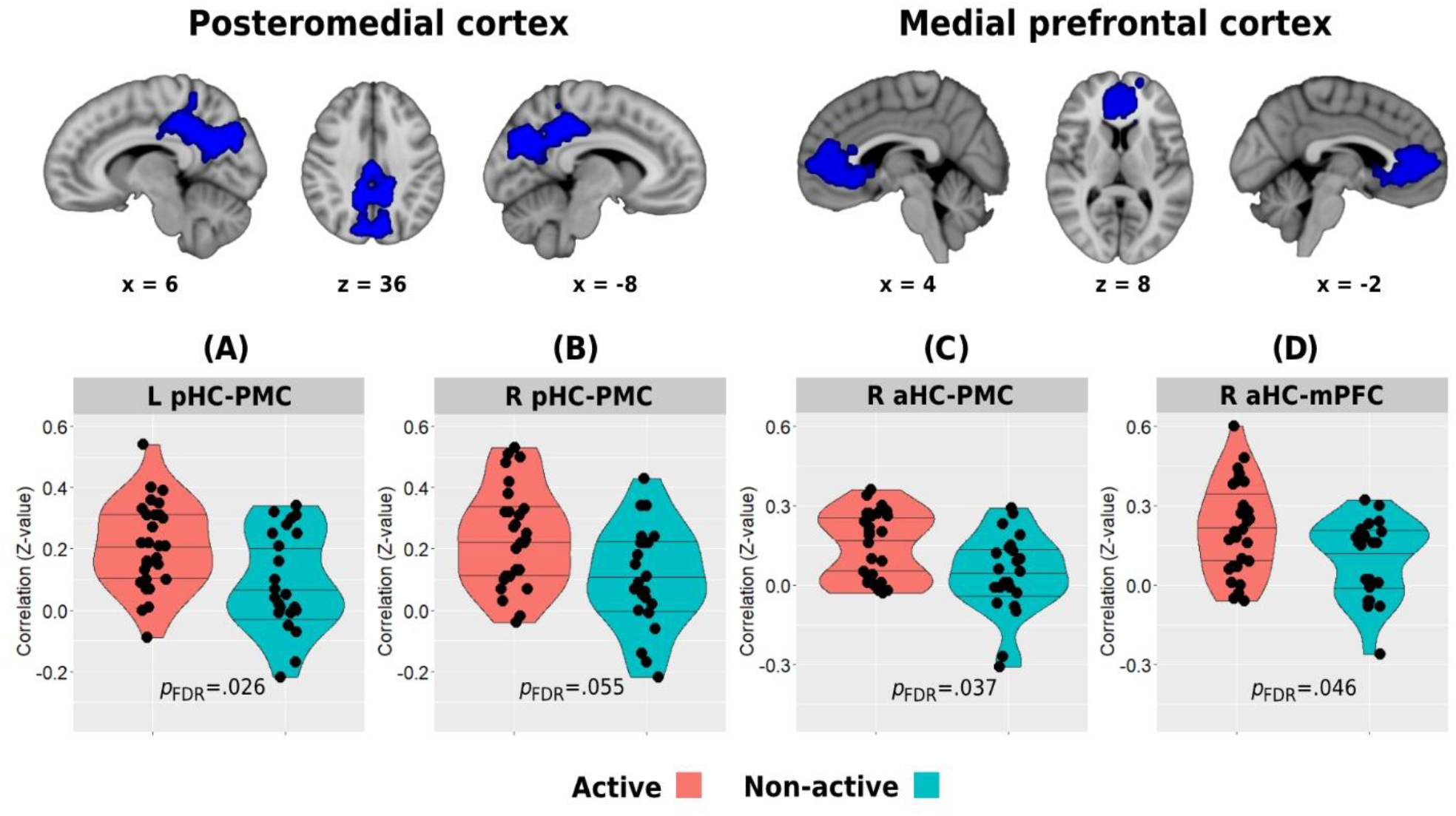
Between-group differences in resting-state functional connectivity of the posterior (A&B) and anterior (C&D) hippocampi with the posteromedial and medial prefrontal cortical clusters which demonstrated decreased activity during the memory encoding paradigm. Top row demonstrates the DMN clusters of interest. Horizontal lines in between-group plots represent 25%, 50%, and 75% quantiles. aHC = anterior hippocampus; L = left; mPFC = medial prefrontal cortex; pHC = posterior hippocampus; PMC = posteromedial cortex; R = right

**Figure 4.**
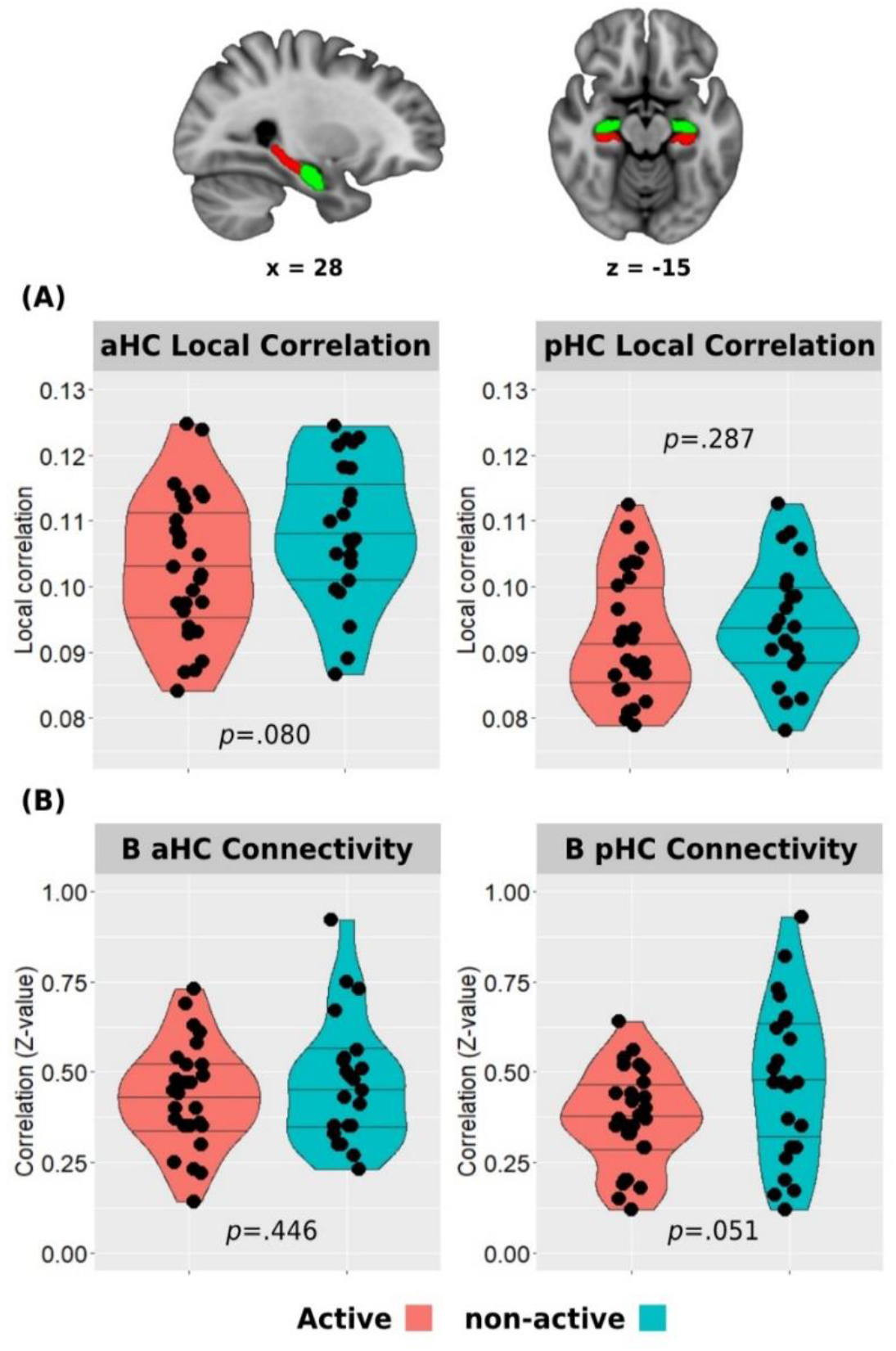
Between-group differences in hippocampal resting state local correlation (A) and between-hemisphere anterior and posterior hippocampal resting state functional connectivity (B). Top row demonstrates the anterior (green) and posterior (red) hippocampal masks used in the analyses. Horizontal lines in between-group plots represent 25%, 50%, and 75% quantiles. aHC = anterior hippocampus; B = bilateral; pHC = posterior hippocampus

### 2.7 General cognitive evaluation

All participants were evaluated for general cognitive functioning and were screened for objective general cognitive decline using the Montreal Cognitive Assessment (MoCA) ^50^.

### 2.8 Statistical analysis

Statistical analyses and visualizations were performed and constructed with IBM SPSS Statistics v24, and R 4.0.3. Between-group differences were tested using non-parametric permutation testing. These procedures were conducted with 10,000 iterations, permuting each hippocampal or memory measure, and preserving the original group sizes. Then, the observed between-group means difference in each measure was compared to all between-group differences obtained from the permuted null distribution. *P*-values were determined as the probability of getting equal or greater between-group difference based on the null distribution. Correlations between MAC and hippocampal and memory measures were evaluated using non-parametric partial Spearman’s rank correlations controlling for age, sex, and years of education. Since the implemented recognition task has no well-established norms, a linear regression was used to create standardized residuals of the memory scores controlled for age, sex, and education, which was used as the memory performance measure for further analyses. In addition, since the local correlation of bilateral anterior or posterior hippocampi were highly correlated between the homologous regions, we used the average local correlation value of the bilateral anterior/posterior hippocampal regions for further analyses. Between-group differences in continuous background characteristics (i.e., age, education, MoCA) were evaluated with Wilcoxon-Mann-Whitney U test, while difference in the proportion between males and females in the groups were assessed using the Chi-squared test. Hippocampal-DMN connectivity between-group tests’ and correlations’ *p*-values (8 tests each) were corrected for multiple comparisons using the False Discovery Rate (FDR) method ^51^.

## 3. Results

### 3.1 Study participants

The active and non-active groups did not differ in any background characteristics, including years of education and general cognitive functioning (*p*>.05) (**Table 1**).

**Table 1.** General socio-demographic and cognitive characteristics of participants (means ± SD).

| Variable | Whole sample<br>(n=52) | Aerobically<br>active<br>(n=29) | Non-active<br>(n=23) | Between-group *p*-value |
| --- | --- | --- | --- | --- |
| Age (y) | 70.83 ± 3.9 | 70.34 ± 4.0 | 71.43 ± 3.7 | .247 |
| Sex | 22 F/30 M | 11 F/18 M | 11 F/12 M | .473 |
| Education (y) | 15.84 ± 3.3 | 16.17 ± 3.7 | 15.41 ± 2.8 | .644 |
| MoCA (raw score) | 24.81 ± 2.5 | 25.13 ± 2.2 | 24.39 ± 2.8 | .349 |

y = years; F = female; M = male; MoCA = Montreal Cognitive Assessment; SD = standard deviation

### 3.2 Aerobic exercise habits and MAC characteristics

The study participants report of weekly aerobic activity ranged from being completely sedentary (not engaging in exercise-oriented activities, n=18) to 6 days per week (n=3). Seventeen participants reported exercising 3 times per week. Exercising on a single day (n=5), twice a week (n=3), 4 times per week (n=4), and 5 times (n=2) were also reported. The active group demonstrated statistically significant higher weekly frequency of exercise sessions and MAC values (**Table 2**).

**Table 2.** Aerobic activity and fitness characteristics of participants (means ± SD).

| Variable | Whole sample<br>($n=52$) | Aerobically<br>active<br>($n=29$) | Non-active<br>($n=23$) | Between-group $p$-value |
| --- | --- | --- | --- | --- |
| Aerobic exercise<br>(d/week) | $2.04 \pm 1.8$ | $3.48 \pm 1.1$ | $.22 \pm .4$ | **<.001** |
| MAC (ml/kg/min) | $25.25 \pm 8.3$ | $30.58 \pm 6.2$ | $18.86 \pm 3.8$ | **<.001** |

d/week = days per week; kg = kilogram; MAC = maximal aerobic capacity; min = minute; ml = milliliter; SD = standard deviation

### 3.3 Hippocampal activation

#### 3.3.1 Between-group differences

Whole brain analysis of all participants revealed bilateral anterior hippocampal activation during the associative memory encoding task (**Figure 2A**). Both groups demonstrated positive mean activation level in both right and left anterior hippocampi during the task. However, the active group activation levels were significantly lower compared to the non-active group (right anterior hippocampus: .09±.31 vs. .38±.31, *p*=.002; left anterior hippocampus: .13±.31 vs. .33±.32, *p*=.027) (**Figure 2B**).

#### 3.3.2 Association with MAC

In accordance with the between-group results, higher MAC was negatively correlated with both right and left anterior hippocampal activation level during the memory encoding task (*r*(38)=-.439, *p*=.004, and *r*(38)=-.334, *p*=.031, respectively) (**Figure 5A&B**).

**Figure 5.**
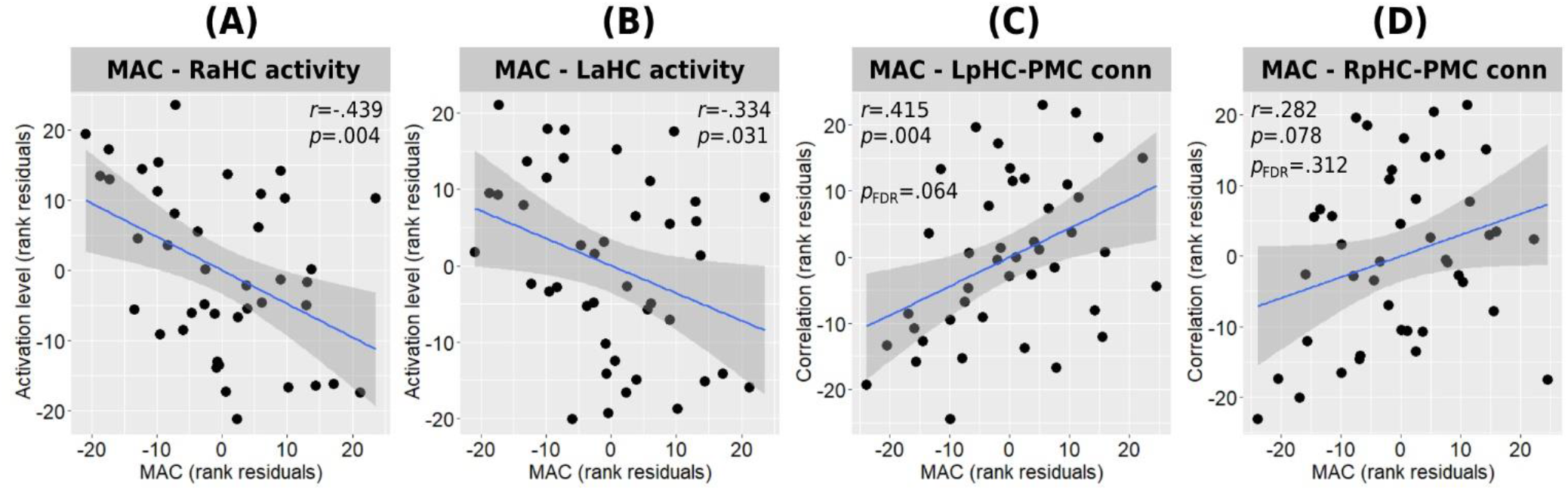
Associations between MAC and hippocampal function. Correlations between MAC and right (A) and left (B) anterior hippocampal activity during memory encoding, and with the connectivity strength of the left (C) and right (D) posterior hippocampi connectivity with the posteromedial cortex are presented. All correlations are adjusted for age, sex, an education. MAC = maximal aerobic capacity; L/RaHC = left/right anterior hippocampus; L/RpHC = left/right posterior hippocampus; PMC = posteromedial cortex

### 3.4 Distant hippocampal functional connectivity

#### 3.4.1 Between-group differences

Whole brain analysis of deactivation contrast during memory encoding revealed several clusters of areas demonstrating decreased activity that are usually attributed to the DMN (see **Table S1**). The two main and larger clusters observed corresponded to the posteromedial and medial prefrontal cortices, which were used as the ROIs for the HC-DMN RSFC analysis (**Figure 3**). In turn, between-group analysis revealed both the left and right posterior hippocampi to demonstrate higher RSFC with the posteromedial cortex in the physically active group compared to the non-active group (left posterior hippocampus: 21±.14 vs. .09±.16, *p*=.009, *p*FDR=.026; right posterior hippocampus: 23±.16 vs. .11±.17, *p*=.007, *p*FDR=.055) (**Figure 3A&B**). In addition, the aerobically active group demonstrated higher RSFC between the right anterior hippocampus and both DMN hubs (posteromedial: .15±.13 vs. .04±.15, *p*=.009, *p*FDR=.037; medial prefrontal: 21±.17 vs. .10±.15, *p*=.023, *p*FDR=.046) (**Figure 3C&D**). All other hippocampal-DMN regional connectivity values were not different between the groups (*p*>.40).

#### 3.4.2 Association with MAC

Higher MAC was found to demonstrate moderate positive correlation with the connectivity strength of the left posterior hippocampus and the posteromedial cortex (*r*(38)=.415, *p*=.004, *p*FDR=.064) (**Figure 5C**) and a weak-to-moderate positive correlation with the connectivity strength of the right posterior hippocampus and the posteromedial cortex which demonstrated a trend toward statistical significance before multiple tests correction (*r*(38)=.282, *p*=.078, *p*FDR=.312) (**Figure 5D**). All other hippocampal-DMN connectivity values demonstrated weak to negligible correlations.

### 3.5 Local hippocampal functional connectivity

#### 3.5.1 Between-group differences

The active group demonstrated lower resting state local correlation in both anterior and posterior hippocampus, but only the anterior hippocampus reached a trend for statistical significance (anterior: .103±.011 vs. .108±.011, *p*=.080; posterior: .092±.010 vs. .094±.009, *p*=.287) (**Figure 4A**). In addition, the bilateral posterior hippocampi resting state connectivity was lower in the active compared to the non-active group (.37±.14 vs. .47±.22, *p*=.051). No difference was found between in the bilateral anterior hippocampus connectivity across the groups (.43±.14 vs. .47±.17, *p*=.446) (**Figure 4B**).

#### 3.5.2 Association with MAC

MAC demonstrated weak correlations with anterior and posterior intra-hippocampal local correlations (*r*(38)=-.232, *p*=.150 for both regions), as well as with inter-hemispheric hippocampal connectivity (anterior: *r*(38)=.089, *p*=.585; posterior: *r*(38)=-.166, *p*=.305).

### 3.6 Memory performance and relationship with hippocampal activation during the task

Between-group analysis revealed a statistically significant difference between the groups with the active group demonstrating higher performance on the memory recognition task compared to the non-active group (residual score .24±.95 vs. -.30±.93, *p*=.043) (**Figure 6A**). In addition, higher MAC was positively associated with higher performance in the recognition task (*r*(39)=.354, *p*=.023) (**Figure 6B**). No correlation was found between anterior hippocampal activity level and memory performance on the whole sample level in both right (Spearman rho(45)=-.139, *p*=.350) and left (Spearman rho(45)=-.199, *p*=.180) hippocampi while controlling for age, sex, and education, or when examined separately within each group for the active (right: Spearman rho(24)=.009, *p*=.967, left: Spearman rho(24)=.009, *p*=.964) and non-active (right: Spearman rho(16)=.156, *p*=.536, left: Spearman rho(16)=-.261, *p*=.296) groups.

**Figure 6.**
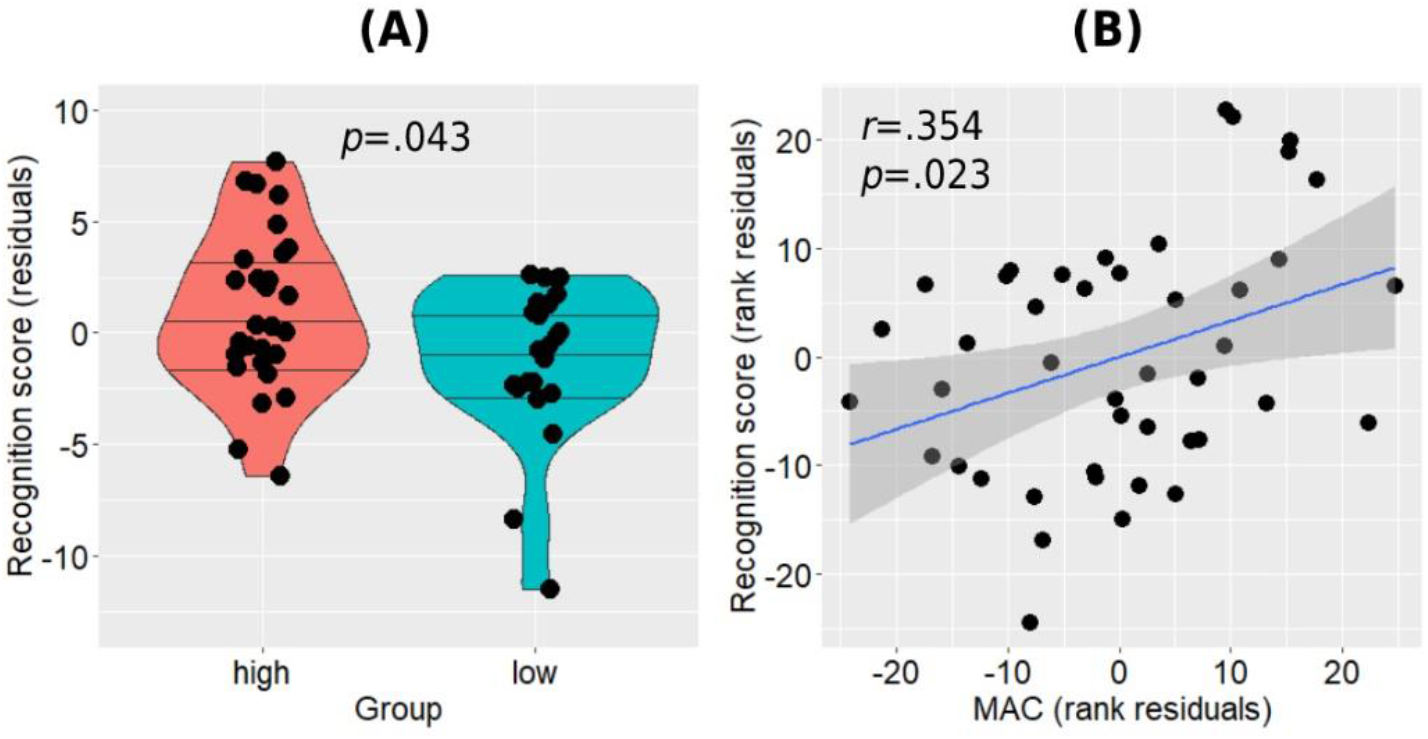
Memory recognition performance differences between the groups (A) and its association with MAC while controlling for age, sex, and education (B). Horizontal lines in between-group plots represent 25%, 50%, and 75% quantiles. MAC = maximal aerobic capacity

## 4. Discussion

As aging has been demonstrated to be associated with compromised brain structure and function ^8,9,52,53^, research has been conducted to investigate lifestyle factors which may attenuate this deterioration with advanced age ^54,55^. Physical exercise, aerobic in particular, has been associated with reduced risk of cognitive decline and dementia ^22^, and hippocampal resilience in older adults ^28,29,56^. However, studies investigating the neuroprotective relationship between physical activity and hippocampus have been mainly focusing on hippocampal structure ^56^ and to a lesser extent on hippocampal cerebrovascular properties ^29,57^. The current study aimed to broaden this line of research by investigating the relationship between aerobically active lifestyle and hippocampal function in cognitively healthy older adults using both resting state and task-based fMRI experiments. In addition, we aimed to examine whether MAC, a potential mediator of aerobic exercise effects may be associated with hippocampal functional characteristics in a dose-response manner.

While anterior hippocampal activity patterns during memory encoding have been generally demonstrated to exhibit age-related decline, some older individuals have been shown to exhibit increased hippocampal activity, or hyperactivity, exceeding values demonstrated in even younger counterparts. Both active and non-active groups in the current study demonstrated increased bilateral anterior hippocampal activation during the associative memory task. However, the non-active group demonstrated significantly higher activity level compared to the active group, with this difference being more pronounced in the right anterior hippocampus. In accordance with this trend, MAC was also negatively correlated with bilateral anterior hippocampi activation level. Mormino et al. ^9^ found successful episodic memory encoding to be associated with right hippocampal hyperactivity in cognitively normal older adults with high Aβ burden compared to young adults and older adults without significant AD pathology. While not reporting on the specific relationship between the hippocampal hyperactivity and post-task memory performance, they demonstrated that the general increase in brain activity observed in the high-risk participants was associated with favourable memory performance, suggesting for a compensatory mechanism as being previously shown in patients with amnestic mild cognitive impairment ^13,58^. This explanation was ruled out in the cognitively healthy sample of the current study since while the active group which demonstrated lower hippocampal activity also showed higher memory performance, increased hippocampal activity was also not associated with better memory performance in each group separately or at the whole sample level. Previous works supported the possibility that increased hippocampal activity may indeed reflect an aberrant dysfunctional activity pattern. Nyberg et al. (2019) ^8^ found older adults exhibiting right anterior hippocampal hyperactivity to demonstrate lower memory performance, higher genetic risk of AD, and higher incidence of deteriorating to dementia at follow-up. Leal et al. (2017) ^59^ found increased right hippocampal activity during memory encoding to be associated with longitudinal accumulation of Aβ several years later, which in turn was associated with steeper memory decline. Importantly, this association was not demonstrated for other cortical areas demonstrating increased activity prior to follow-up. Hippocampal hyperactivity during different types of memory encoding tasks was also shown to be associated with increased tau pathology and lower memory performance in cognitively normal older adults ^60,61^. Further evidence supporting the potential link between AD pathology and hippocampal hyperactivity come from findings in genetic models of the disease. Aberrant hyper-excitatory activity in the hippocampus is a consistent finding in animal models of AD ^62,63^, whereas young adults carrying the mutation for familial subtype of AD have been repeatedly demonstrating right anterior hippocampal hyperactivity during memory encoding compared to non-carriers controls ^64,65^. Furthermore, increased hippocampal activity during memory encoding has also been shown in young individuals at risk for late onset sporadic AD, i.e., APOE ε4 carriers ^66,67^. The observed relationship between hippocampal hyperactivity and Aβ accumulation may be a reflection of a vicious neuropathological cycle in which over-excitatory neurons drive local Aβ aggregation, which in turn further increases neuronal excitability levels ^62,68^. While aerobic exercise has been repeatedly demonstrated to be associated with lower risk of AD and dementia ^20,22,69^, a mechanism which may link aerobic exercise with hippocampal hyper-excitability and dysfunction attenuation in aging may lie in Aβ metabolism. AD model mice demonstrated lower hippocampal amyloid plaque load following exercise intervention, with greater training intensity resulting in lower Aβ burden ^70^. Furthermore, evidence for increased glymphatic clearance of hippocampal amyloid plaques in aged mice was also demonstrated following a single session of aerobic exercise ^71^. One study in already diagnosed human patients with AD did not find aerobic intervention to affect Aβ levels, however the late clinical stage of the participants on the AD continuum, the relatively short duration of the intervention, and significantly higher Aβ levels in the exercise group at baseline may account for the lack of interventional efficacy observed in the study ^72^. In addition, while normal hippocampal function requires a balance between excitation and inhibition, abnormal hippocampal excitation in pathological aging has been also linked to GABAergic inhibitory dysfunction ^73,74^. Aerobic exercise in turn, has been demonstrated to induce neuroprotective inhibitory modulation in the hippocampus of hyper-excitatory epileptic rats ^75,76^. This in turn, suggests another potential neurobiological mechanism which may underlie the relationship between aerobic exercise and attenuated hippocampal hyperactivity observed in the current study.

The second primary finding in our work are the differences observed in hippocampal resting state connectivity between the groups. Namely, aerobically active lifestyle (and to a lesser extent MAC) was found to be associated with higher distant hippocampal functional connectivity with core hubs of the DMN, i.e., the posteromedial and medial prefrontal cortices, as well as with lower local intra-hippocampal connectivity. The hippocampus is a part of the medial temporal subsystem of the DMN ^16^, and these regions have been shown to functionally cooperate in cortico-hippocampal memory networks ^5,6^. Previous works demonstrated decreased cortico-hippocampal connectivity in healthy older adults. While Damoiseaux et al. (2016) ^18^ found only the posterior hippocampus to demonstrate age-related decline in functional connectivity with the DMN, Salami et al. (2014) ^19^ demonstrated this age-related effect to occur in both anterior and posterior hippocampal subparts, and that higher connectivity values were positively correlated with episodic memory performance. We found a link between physically active lifestyle and higher bilateral posterior hippocampi connectivity with the posteromedial cortex while also higher connectivity of the right anterior hippocampus with both posteromedial and medial prefrontal cortices. In addition to higher distant connectivity, the active group in our study also demonstrated a trend toward lower intra-hippocampal connectivity expressed as lower local correlations within the anterior hippocampus and between the posterior hippocampi across the two hemispheres. Higher within- and between-hippocampi correlations have been previously demonstrated with increasing age, negatively correlating with memory performance and load of AD pathology ^17,19^. Harrison et al. (2019) ^17^ further demonstrated that increased local within hippocampal correlation, represented by higher regional homogeneity, was associated with decreased cortico-hippocampal connectivity, suggesting that both age and AD pathology may be associated with increasing hippocampal functional disconnection and isolation. In contrast, our findings provide evidence that aerobic activity and physically active lifestyle may constitute a neuroprotective factor in face of this aspect of hippocampal dysfunction. In turn, these results extend previous works which demonstrated a relationship between physical activity and increased DMN connectivity ^77,78^. Reduced functional connectivity of the DMN is associated not only with aging ^19^ but also with early biomarkers of AD pathology, as being highly sensitive regions to early Aβ accumulation ^79^. Given the evidence pointing at the potential role aerobic activity may have in increasing of the glymphatic removal of Aβ deposition from the interstitial space, it may underlie, at least partially, the observed relationship with higher functional connectivity of the two core hubs of DMN with the hippocampus. In addition, one of the hallmarks of exercise-induced mechanisms observed in animal models is the up-regulation of brain and hippocampal brain-derived neurotrophic factor (BDNF) ^26,80,81^. Among its versatile functions, BDNF promotes neurite outgrowth and synaptogenesis, plays a significant role in synaptic plasticity, and is involved in learning and memory processes ^82,83^. In human older adults, increased serum levels of BDNF was associated with increased parahippocampal functional connectivity following a year of aerobic intervention, supporting the potential role of BDNF as a neurobiological mediator of exercise-induced changes in brain function ^84^. It is important to note that although MAC was associated with some of the hippocampal measures we examined in the current study, it did not explain all the between-group differences observed, especially in terms of distant and local hippocampal connectivity. Although it may support a potential mediating role of MAC in some of the exercise-related hippocampal function effects, it also suggests that other potential mechanisms play a role in mediating this relationship.

### 4.1 Study limitations

The main limitation of the current study lies in its cross-sectional nature. Although previous evidence from animal models and human participants support the observed relationship between aerobic exercise and hippocampal function from a neurobiological mechanistic point of view, we cannot state that physically active lifestyle is indeed the causal mediator of the attenuation in hippocampal dysfunction observed in the current sample of cognitively healthy older adults. Instead, these results should serve as a starting point for future longitudinal and interventional studies that are needed in order to explore the time-dependency between aerobic exercise and functional adaptations of the hippocampus in human aging and to establish a causal effect.

## 5. Conclusions

While the relationship between physical activity and hippocampal function in human aging has been sparsely investigated, the current study provides evidence for a potential relationship between lifestyle habits of aerobic exercise and attenuation of different aspect of hippocampal dysfunction, normally observed in older individuals, namely alteration in patterns of hippocampal activity and connectivity. By that the current research extend previous works demonstrating the relationship between physical activity and aging brain in general, and specifically the neuroprotective effect of aerobic exercise on the aged hippocampus. In addition, our results suggest that although MAC may be associated with some aspects of hippocampal function in older adults, other factors may also contribute to the relationship between aerobic exercise and distinct hippocampal functions.

## Acknowledgements

We thank Prof. Dafna Ben Bashat for her consultation during the research process. We thank Dr. Moran Artzi for help with planning of the acquisition sequences, and Prof. Yuval Nir, for his thoughtful advice. We also thank Dr. Avraham Man for his assistance with conducting the cardiopulmonary testing.

---

**Table S1.** Significant clusters demonstrating deactivation during memory encoding (cluster-corrected).

| Cluster | *x* | *y* | *z* | Cluster Size | z-value max |
| --- | --- | --- | --- | --- | --- |
| 1 | 4 | -30 | 44 | 4308 | 5.26 |
| 2 | 0 | 42 | -4 | 2701 | 4.77 |
| 3 | -40 | -74 | 40 | 1523 | 4.86 |
| 4 | 66 | -20 | -4 | 1276 | 4.91 |
| 5 | 46 | -74 | 38 | 960 | 4.73 |
| 6 | 34 | -52 | 4 | 682 | 5.06 |
| 7 | -30 | -58 | 4 | 568 | 4.73 |

Coordinates are in MNI space

